# L2 loop engineering enhances the enzymatic activity and synergism property of the AA16 family Lytic polysaccharide monooxygenase enzyme from *Aspergillus fumigatus*

**DOI:** 10.1101/2025.08.27.672579

**Authors:** Musaddique Hossain, Shalini Das, Soumyadeep Ray, Nilanjana Bose, Bishwajit Kundu, Sudit S. Mukhopadhyay

**Affiliations:** Department of Biotechnology, National Institute of Technology, Durgapur 713209, India; Department of Computational Biology, Indraprastha Institute of Information Technology, Delhi, Okhla Phase III, New Delhi, 110020, India; Kusuma School of Biological Sciences, Indian Institute of Technology Delhi, Hauzkhas 110016

**Keywords:** Protein engineering, LPMO, cellulose degradation, L2 Loop, Cellulase

## Abstract

Lytic polysaccharide monooxygenase (LPMO) is an enzyme that has enormous potential for industrial applications. It has a synergistic effect with the cellulase enzyme complex. The LPMO enzymes typically adopt a compact β-sandwich fold that consists of a 7 to 9 β-strand with a flat active site containing copper in its active site. There are a few loops in LPMO, among them the L2 loop is reported to take a key role in shaping the active site and substrate binding. Now, in this work, we want to investigate the role of the L2 loop in enzymatic activity and synergistic effect. In achieving our goal, we have replaced the L2 loop of our concerning *Af*LPMO16 with other L2 loops from different LPMOs: *Hi*LPMO9B (PDB: 5NNS), *Mc*LPMO9 (PDB: 7NTL) and *Cs*LPMO9 (PDB: 7EXK). Interestingly, L2 loop replacement from *Cs*LPMO9 (PDB: 7EXK) showed enhanced activity and synergism compared to others. The secondary structural analysis by circular dichroism also suggested that it changed the structure significantly. Moreover, this is the first report of complete L2 loop engineering in LPMO.

## Introduction

Lytic polysaccharide monooxygenase (LPMO) is an enzyme that has enormous potential for industrial applications (1). This synergistic effect with cellulase and other polysaccharide-degrading enzymes is a crucial aspect for industrial limelight. The structural architecture of the typical LPMO was elucidated in several studies (2–4). The main structural architecture suggests that it has a copper-containing exposed active site. The copper is coordinated mainly by two histidine braces (5). Other than the active site, the enzyme has a distorted fibronectin or immunoglobulin-like β-sandwich fold. It generally contains two β-sheets that consist of seven to nine β-strands (6–8). These β-strands form a compact pyramidal core that stabilises the whole structure of the enzyme LPMO. These strands are connected by some loops and helices. There are mainly five loops in LPMO cited in different articles. The five loops are L2, L3, LS (short loop), L8 and LC (long loop) (9). These loops are named based on their position in the crystal structure of LPMOs, such as the L2 loop positioned between β-strand 1 and β-strand 2 in AA9 LPMOs and in AA10 LPMOs, it is positioned between β-strand 1 and β-strand 3. Whereas the L3 loop is positioned between β-strands three to four. The LS loop is a short loop just opposite to the L2 loop between specific β-strands which involved in substrate binding (10). The L8 loop is positioned between β-8 to β-9. The LC loop is a long C-terminal loop (9).

Among these loops, L2 is the crucial one due to its involvement in substrate binding and determination of substrate specificity and stability (9). The first report of a tryptophan residue (W57) crucial for substrate bonding was an indication that this stretch of amino acids is involved in substrate binding (11). The other amino acids, like Tyr-54, Glu-55, and Glu-60, present in the L2 loop are crucial for chitin (substrate) binding (12). Another protein engineering study showed the aromatic replacement (Y56W) did not alter the detectable substrate binding but preferential synergistic effect (13). Multiple single-mutation studies revealed that the residues in the L2 loop are crucial for activity, also beyond the substrate binding (14). All these studies confirm that the various amino acids, aromatic and polar residues in the L2 loop have an important role in binding with specific substrates, also they can influence the catalytic activity (15).

All these studies mainly focused on a single mutation and specific residues. But there is no such protein engineering approach that has been considered for the whole L2 loop. There is a scope to study the complete L2 loop or the replacement of the L2 loop and its effect on substrate binding and affinity, enzymatic activity and protein stability. However few questions are integrated with the L2 loop replacement. First, does the substrate specificity completely or partially depend on the L2 loop? Second, does the regioselectivity depend on the L2 loop? Third, is there any role in substrate profiling, catalytic activity and synergism? To answer these questions, we replace the L2 loop of *Af*LPMO16 with three different L2 loops for three different LPMOs from the AA9 family. The three L2 loops are taken from *Hi*LPMO9B (PDB: 5NNS) (16), *Mc*LPMO9 (PDB: 7NTL) (17) and *Cs*LPMO9 (PDB: 7EXK) (9). These three LPMOs have distinct structural features; the L2 loop of *Hi*LPMO9B does not have any helix, whereas the L2 loop of *Mc*LPMO9 has two short helices or a fragmented helix, and the L2 loop of *Cs*LPMO9 has a single helix. The comparison of the enzymatic activity revealed that the *Cs*LPMO9 showed the highest reported activity and synergism (9). Our focus is to check the effect of the replacement of the L2 loops with the native L2 loop of *Af*LPMO16. The effect means starting from the structural integrity of the chimeric protein to the enzymatic activity. We have successfully developed three mutant proteins with the replaced L2 loops: first, *Af*LPMO_5NNS; the altered L2 was taken from *Hi*LPMO9B, second, *Af*LPMO_7NTL, and third, *Af*LPMO_7EXK, where the L2 loop was altered from *Mc*LPMO9 and *Cs*LPMO9, respectively. After the successful mutation generation, we confirmed structural integrity by circular dichroism and checked the different aspects of enzymatic activity. Interestingly, we found that the replacement of the L2 loop can influence the total enzymatic activity, even though the L2 loop can only be the signature of the enzymatic activity of LPMO. Here in this study, we first design the sequence chimeric proteins *Af*LPMO_5NNS, *Af*LPMO_7NTL and *Af*LPMO_7EXK by only replacing the L2 loops. Then we predicted the three-dimensional structure, and using that structure, we performed the MD simulation and docking. Then we cloned the mutant proteins and successfully expressed them in *Pichia pastoris* (18). We performed biophysical studies to check the physicochemical properties and structural integrity. Lastly, we performed the enzyme assays with different substrates and compared the synergistic effect of the mutant enzymes with the native *Af*LPMO16. Here, we first report the successful loop engineering in LPMO (AA16) and the role of the specific L2 loop in controlling the overall enzymatic activity and synergism.

## Materials and methods

### Sequence and design of the loop-altered constructs and three-dimensional structure prediction

The native LPMO from *Aspergillus fumigatus Af*LPMO16 was cloned and the protein sequence was taken, and a three-dimensional model structure was created using Boltz-1x in the Tamarind Bio website with [Cu^2+^] as smiles (https://app.tamarind.bio/boltz) (19). This tool is similar to AlphaFold3 and uses the same algorithm (20). We obtained the PDB files and renamed the copper residue name to CU, and renamed the chain of the copper residue from B to A, and modified the residue number as (last residue number of protein chain A + 1). After this step, we used PyMol visualizer to visualise the model structure (21). The β-1 and β-2 marked and found the L2 loop and its sequence. Similar steps are followed to find out the L2 loop sequence of the *Hi*LPMO9B (PDB: 5NNS), *Mc*LPMO9 (PDB: 7NTL) and *Cs*LPMO9 (PDB: 7EXK). The corresponding L2 loop sequences are replaced with the native L2 loop sequence to generate three chimeric proteins: *Af*LPMO_5NNS, *Af*LPMO_7NTL and *Af*LPMO_7EXK (table S1). The steps are followed to predict the model structure of the loop-engineered proteins: *Af*LPMO_5NNS, *Af*LPMO_7NTL and *Af*LPMO_7EXK.

### Molecular Dynamics simulation

Molecular dynamics simulation is a challenging job for metalloproteins (22). We developed a new method to run the MD simulation in GROMACS (23). The topology file was created using AMBER (24) then it was converted to gromacs file using code1 (Table S2). Then we used OpenMM – CUDA accelerated to carry out MD simulations, 1fs step size, temp 300K, friction=1 ps (25). First, the AMBER topology (prmtop) is parsed to locate the copper ion (Cu) and the specific coordinating atoms: the backbone amide nitrogen (N) and imidazole ε_nitrogen (NE2) of histidine at position X, the NE2 of histidine at position Y and the hydroxyl oxygen (OH) of tyrosine at position Z. Once the indices are known, the corresponding Cartesian coordinates are read from the coordinate file (inpcrd). Each Cu–ligand pair’s interatomic distance is then computed in Angstroms and converted into nanometers; these distances serve as the equilibrium bond lengths (r_).

A single Custom Bond Force is created to apply harmonic restraints of the form U(r) = 1/2k(r-r_0_)^2^ Where the force constant k is set to 10 000 kJ·mol□^1^·nm□^2^ and each bond has its own r_0_ (the measured Cu–ligand distance). Four bonds are then added to this force: between Cu and HIS X N, Cu and HIS X NE2, Cu and HIS Y NE2, and Cu and TYR Z OH. Finally, this custom bond force is attached to the system so that, during dynamics, the copper ion remains stably coordinated to those four atoms. Then did energy minimisation (5000steps) followed by NVT (50000 steps) (300K) (maxwell-boltzmann velocities) and then NPT (1atm) (300K) (50000steps) {monte-carlo barostat}. Finally, the MD run, Production Run-100 million steps, i.e 100ns.

### Quantum chemical methods

The single-point energy calculation was carried out using ORCA (26). First, all residues within 3.5 Å of the copper atom were selected to define its coordination sphere. The two nearest water molecules were included explicitly to capture the first solvation shell. The first energy minimum from the MD trajectory was used to study the most stable conformer. The final MD frame was analysed to incorporate a realistic snapshot of protein flexibility and solvation. The UKS B3LYP functional was chosen for its proven accuracy with transition-metal centres in ORCA calculations. The def2-TZVPP basis set was used to provide triple-ζ quality and polarisation functions around the copper site. Grimme’s D4 dispersion correction was added to account for non-covalent interactions with surrounding residues. Tight SCF convergence criteria and SlowConv settings were applied to ensure stable SCF convergence in this open-shell system. PAL8 parallelization directives enabled efficient use of multiple CPU cores. A total charge of +2 for the system, a multiplicity of 2 was assigned to represent the Cu(II) (except for *Af*LPMO_7NTL +1 charge and multiplicity 2)(27).

### Cloning and expression

The native *Af*LPMO16 enzyme was initially cloned and sequenced (28). Further, the same sequence was synthesised and cloned by GenScript with an upstream XhoI site for native N-terminal expression in the pPICZαA vector. The chimeric proteins *Af*LPMO_5NNS, *Af*LPMO_7NTL and *Af*LPMO_7EXK were constructed by GenScript; the sequence details are given in the supplementary section (Figure S1). After successful construction of the engineered protein, we linearised the cloned plasmid and transformed it into *Pichia pastoris* X33 cells (29). The positive transformants were selected using Zeocin antibiotic as a selection marker. Then the recombinant proteins were expressed in BMGY media, followed by BMMY media (30). Then, recombinant proteins were purified using His-tag-based affinity chromatography (31). The purified proteins were analysed using 12% SDS-PAGE. The purified protein was further confirmed by western blot analysis using C-Myc antibody and Anti-His antibody (ThermoFisher MA 1-980). The purified proteins are further used for enzymatic analysis.

### Enzyme assays

There are three types of enzyme assays for LPMOs first one is the colourimetric assay using 2,6-DMP as a substrate, the second is the cellulose degradation assay, and the third one is the Amplex red assay, which produces hydrogen peroxide. We performed all of them to determine the enzymatic activity of the *Af*LPMO16 and loop-altered chimeric LPMOs. But there is a step of LPMO activation before going into the details of the assays. LPMO is copper copper-dependent enzyme. During purification, it may lose the copper ion in its active site so we used a 1M CuSO4 solution and mixed it with a 1:1 volume ratio for 1hour at 16°C. The excess CuSO4 was removed using a Zeba protein desalting spin column (ThermoFisher). There were two negative controls in each reaction; one was without copper saturation, another is without enzyme.

### 2,6-Dimethoxyphenol assay

At first, we prepare a stock of 10 mM 2,6-dimethoxyphenol (2,6-DMP) and 5 mM H_2_O_2_(freshly prepared). Then we set the reaction, the total reaction volume was 1ml, and the final concentration of the reactants is 100mM sodium phosphate buffer, pH 6.0, 1mM 2,6-DMP, 100uM H_2_O_2_. 200ug LPMO enzyme. Reaction was carried out for 30minutes at 37°C, and the change in absorbance was measured at 469nm wavelength of electromagnetic radiation (29, 32).

### Amplex red assay

The total reaction volume for this assay is 200 μL. The final concentration of the reactants in the reaction is 100mM sodium phosphate buffer, pH 6.0, 50 μg LPMO and 30 μM ascorbic acid. We followed the protocol mentioned in the Amplex™ Red Hydrogen Peroxide/Peroxidase Assay Kit (Invitrogen)(33). The absorbance was taken at a 560nm wavelength. By using a hydrogen peroxide standard curve, we can calculate the amount of generated H_2_O_2_. Here, we don’t add any substrate so that the hydrogen peroxide released by the uncoupling reaction can be calculated to correlate with the enzymatic activity (34).

### Cellulose degradation assay

We used 1% Avicel®PH-101 (SIGMA) (crystalline cellulose) as a substrate. The final concentration of the reactants in the reaction was: 50mM sodium phosphate buffer, pH 6, 30 μM ascorbic acid, 0.5mg/mL Cellulase (MP Biomedicals LLC) and 200 μg purified LPMO. The reaction took place at 37°C and 50rmp shaker. We measured the amount of D-glucose released in the reaction at different time points using the D-glucose assay kit (Megazyme). For each D-glucose estimation reaction, we used 100uL reaction solution. The solution was centrifuged at 10000 rpm for 10 minutes, and the clear supernatant was taken for the assay, where the soluble D-glucose was estimated.

Cellulose and lignocellulose (alkaline pre-treated raw rice straw) (35) were used to determine the cellulose hydrolysis-enhancing capacity. Rice straw was pre-treated with 5% NaOH (1:10 W/V ratio) at 120□C at 15Psi pressure for 1 hour, and sodium azide (20%) 10μl (per 10ml) was added to the reaction mixture to prevent any microbial contamination. The reaction was performed at 37□C, and the amount of reducing sugar was quantified after 6hours, 24hours and 48hours. Reaction sets were prepared using only cellulase, only *Af*LPMO16, combined *Af*LPMO16 (and mutants) and cellulase and lastly, cellulase with inactivated *Af*LPMO16 (without copper saturation).

### Circular dichroism

The CD spectra were recorded at 25□C temperature on a Jasco J-815 Spectropolarimeter controlled with J-800 software. The spectra were obtained at a 0.1 mg/mL protein concentration in a 50 mM phosphate buffer, pH 7.8, using a 1 mm path length. The data was recorded in triplicate using a wavelength from 250 to 200 nm at a scan rate of 50 nm/min using a 2.0-sec response, where the sensitivity standard was set by the software. The data pitch was 1nm with a 2nm bandwidth. The data measures circular dichroism along the Y-axis in mdeg and the X-axis wavelength in nm. We have also measured High tension voltage (HT) along the Y-axis. CD data was analysed by BEST SEL’s online tool (http://bestsel.elte.hu/).

## Results

### Modelling chimeric *Af*LPMO16

The predicted three-dimensional structure of *Af*LPMO16 satisfies the typical structural feature of the active centre with a Copper ion and the metal ion stabilised by two histidine residues and a Tyrosine residue with two water molecules in its close proximity within 3Å (Fig. 1a). The L2 loop of *Af*LPMO16 is long and contains three short helices. In contrast, the structures of the L2 loops displays distinct features such as the L2 loop from *Hi*LPMO9B (PDB: 5NNS) shows no helical content, *Mc*LPMO9 (PDB: 7NTL) shows two short helices and *Cs*LPMO9 (PDB: 7EXK) shows single helix (Fig. 1b). The predicted structures of loop altered LPMOs also show a stable structure (Figure 1c). The predicted structure shows a decrease in helical structure in *Af*LPMO_5NNS and *Af*LPMO_7NTL.

**Figure 1.**
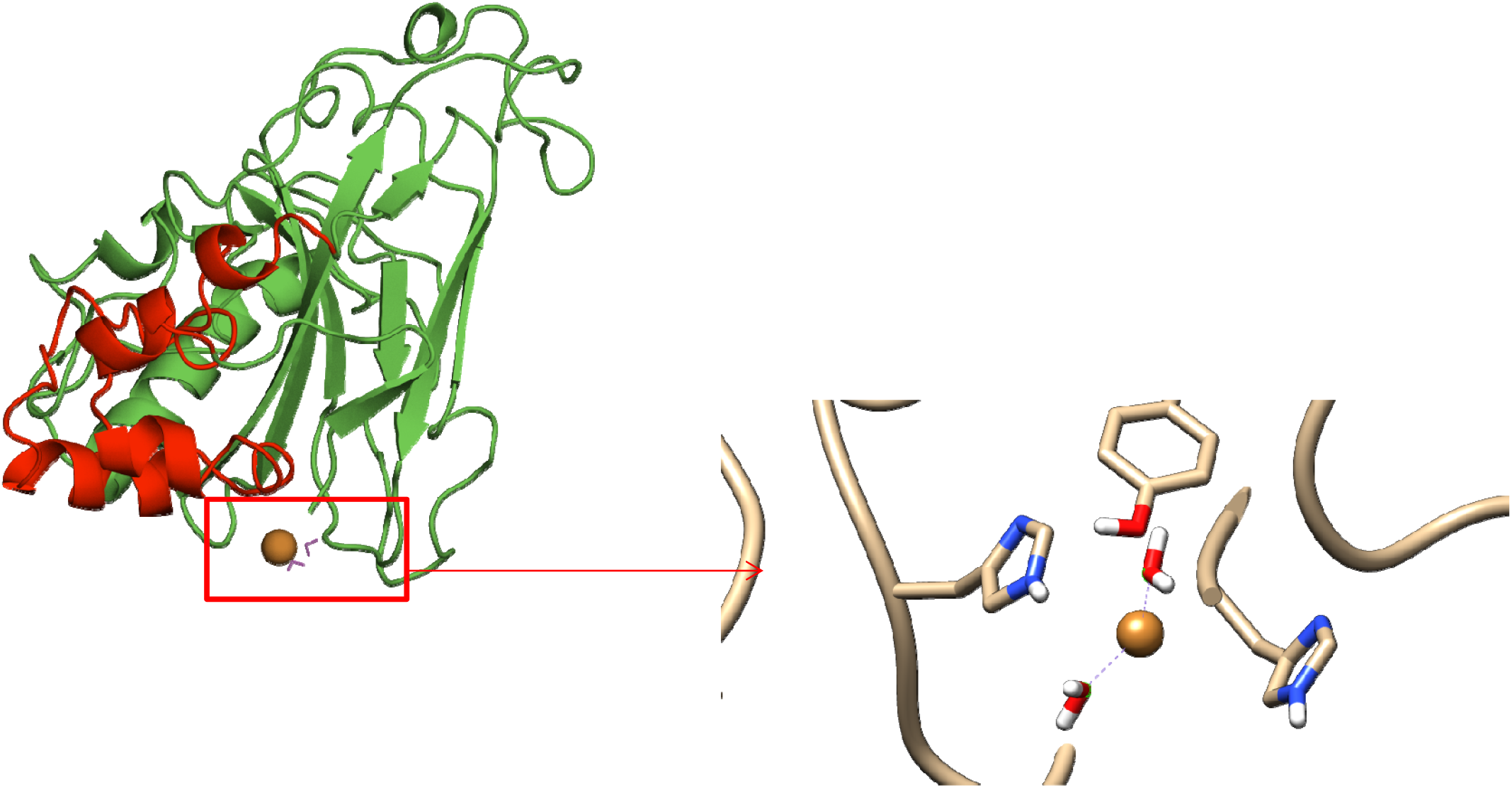
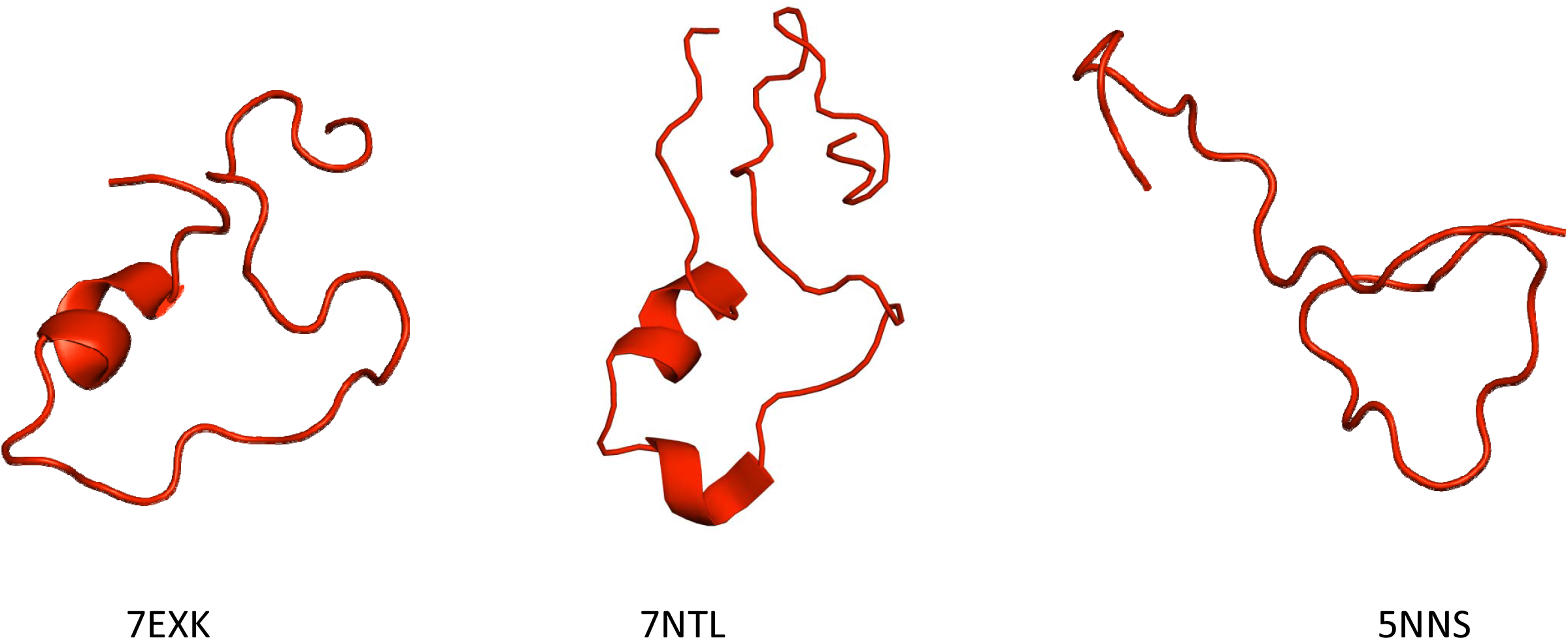
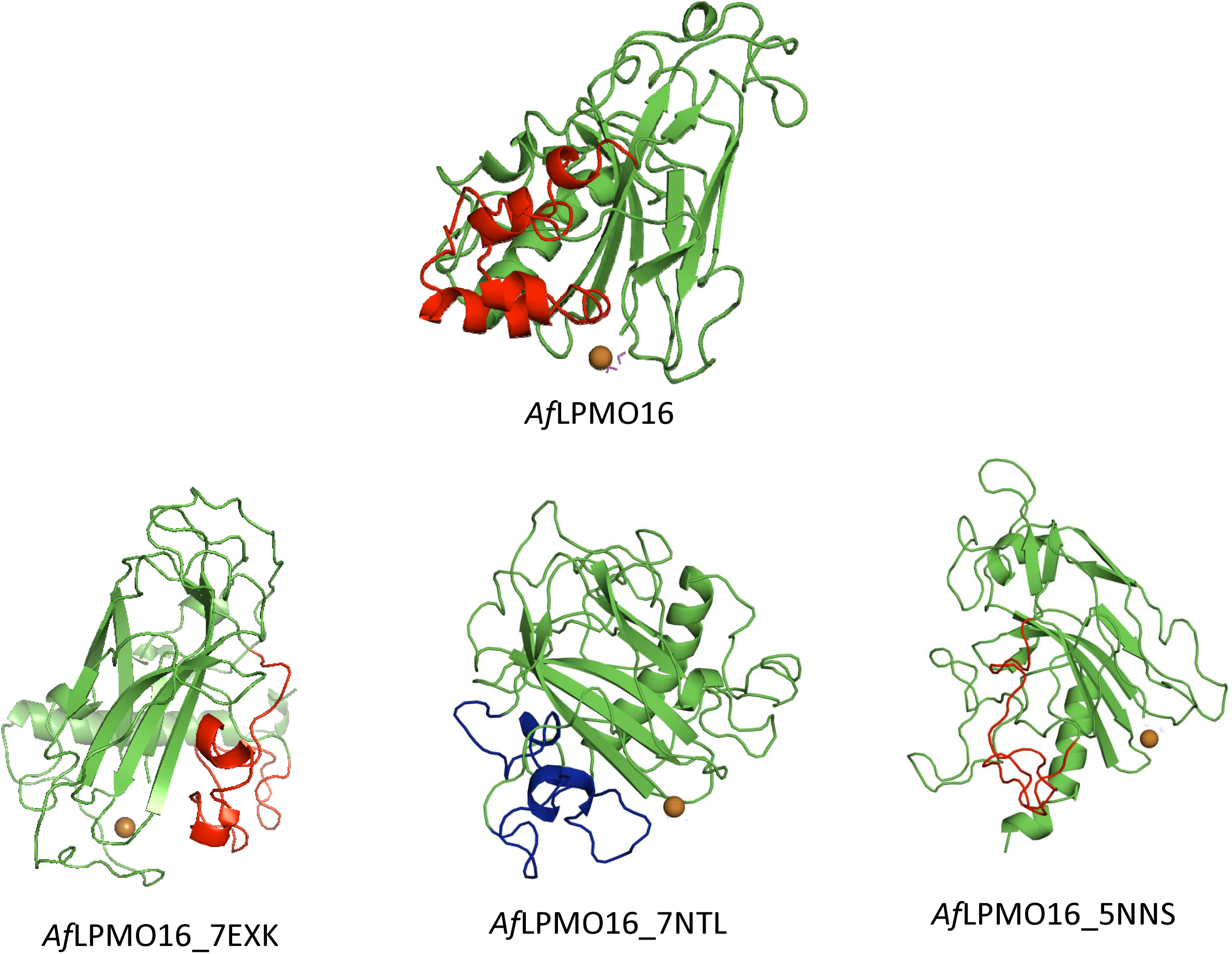
(a) Predicted three dimensional structure of *Af*LPMO16, L2 loop marked in red and catalytic centre zoomed view with His-braces and two water molecule and a tyrosine residue. (b) Structure of three different L2 loop *Hi*LPMO9B (PDB: 5NNS), *Mc*LPMO9 (PDB: 7NTL) and *Cs*LPMO9 (PDB: 7EXK). (c) Model structure of chimeric protein *Af*LPMO_5NNS, *Af*LPMO_7NTL and *Af*LPMO_7EXK.

### Molecular dynamics simulation of Mutant proteins

Structures were further validated by running MD simulation over a period of 100000ps (100ns). The different plots from the trajectory files show the overall stability of the predicted structures. The root mean square deviation plot with time, indicating 2-4A RMSD value for both the *Af*LPMO16 and *Af*LPMO_7EXK, showed an average RMSD of 3Å with respect to C_α_ atom (Fig. 2a). Root mean square fluctuation value versus residues plot suggests the highest fluctuation of L2 loop residues of the *Af*LPMO_7EXK, even the highest fluctuation observed in LC region. Whereas the *Af*LPMO_7NTL shows a lower RMS value compared to the wild type and other mutants (Fig. 2b). This result suggests that the *Af*LPMO_7EXK has higher flexibility compared to the wild type enzyme and other mutant enzymes. Another parameter that indicates the stability is the solvent accessible surface area (SASA). The SASA vs time plot clearly suggests the *Af*LPMO_7EXK has a lower surface area value over the 100ns time period. The range of the SASA is within 100Å^2^ for *Af*LPMO_7EXK enzyme where whereas the *Af*LPMO_5NNS and *Af*LPMO_7NTL have a higher fluctuation rate.

**Figure 2.**
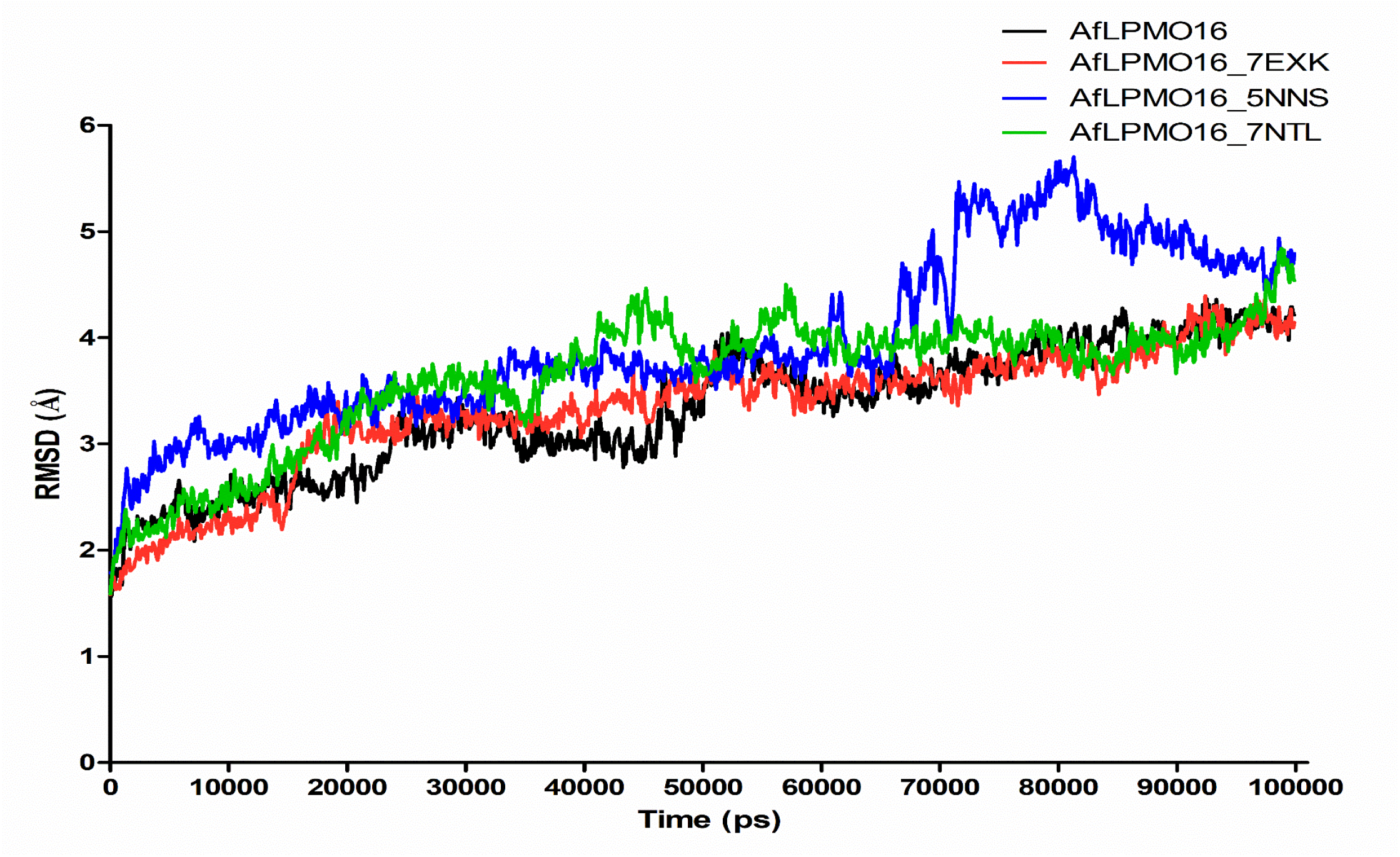
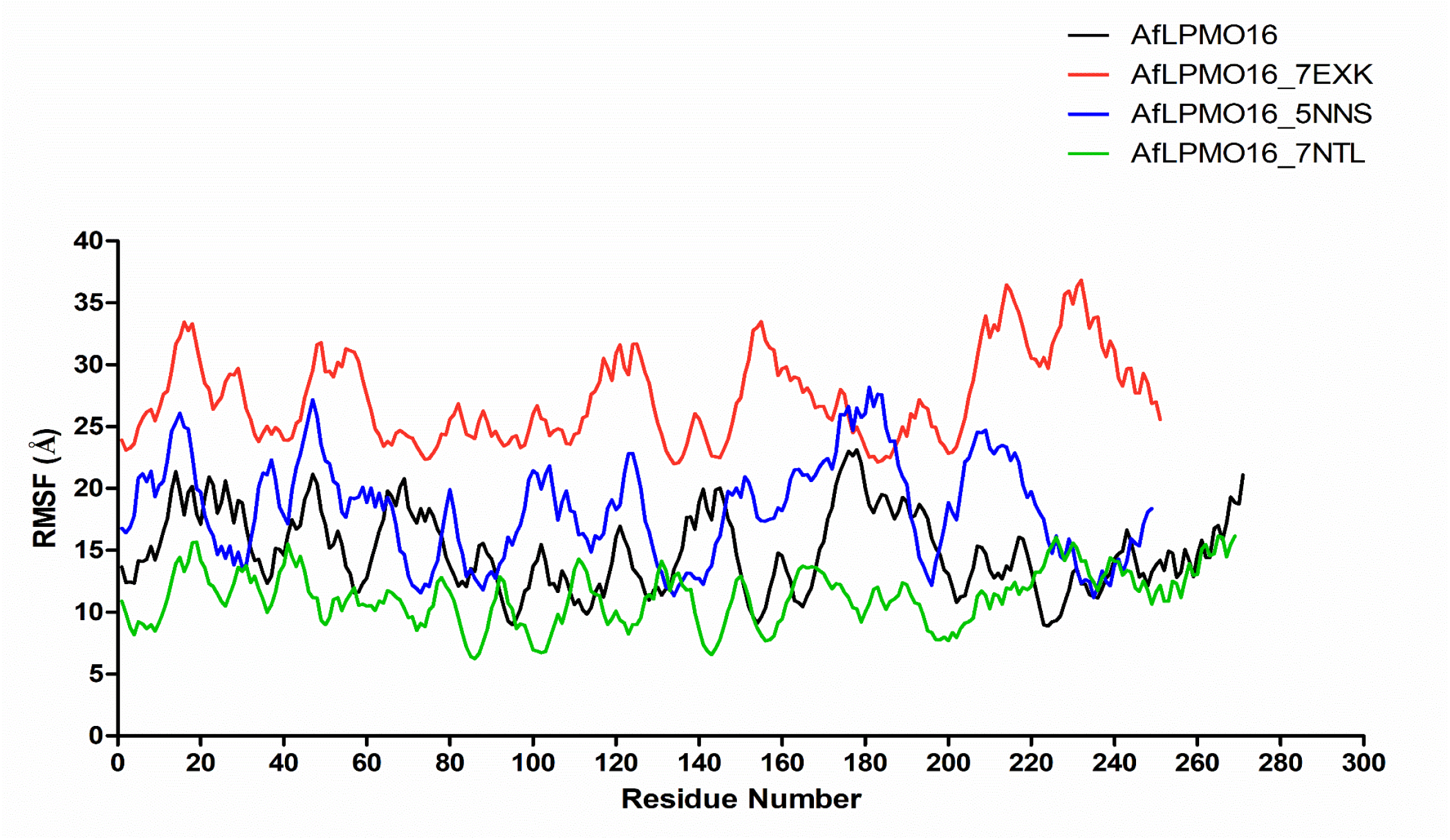
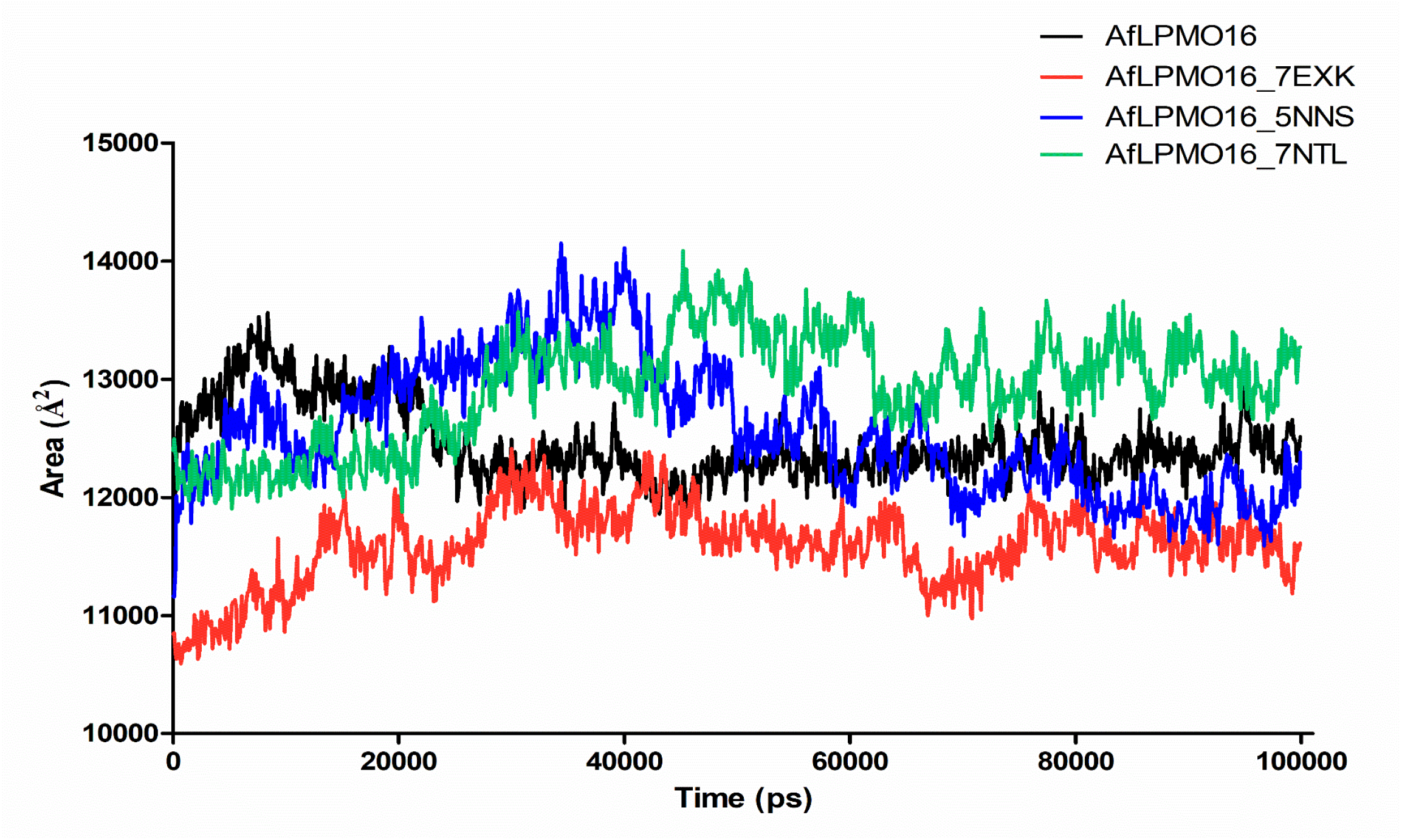
(a) Plot of Root mean square deviation vs Time of *Af*LPMO16 and chimeric proteins *Af*LPMO_5NNS, *Af*LPMO_7NTL and *Af*LPMO_7EXK. (b) Plot of root mean square fluctuation vs residue number *Af*LPMO16 and chimeric proteins *Af*LPMO_5NNS, *Af*LPMO_7NTL and *Af*LPMO_7EXK. (c) Plot of solvent accessible surface area in Å^2^ vs time in Pico-second of *Af*LPMO16 and chimeric proteins *Af*LPMO_5NNS, *Af*LPMO_7NTL and *Af*LPMO_7EXK.

### Quantum mechanical observation of the active site

The electron density contour plot is actually a visual representation of the electronic environment. Here, the four panels represent the electron density contour plot for *Af*LPMO16 and *Af*LPMO_7EXK (Fig. 3). Each of 1st minimum energy state and the last frame is depicted. First minimum energy state of the *Af*LPMO16 has the connected electron density, which confirms the bonds between the copper ion and the residues near it, His1, His and tyrosine. Whereas the first minimum energy state plot of the *Af*LPMO_7EXK. Shows the involvement of two water molecules; the electron density of those molecules is almost connected with the copper ion. Another interesting observation, in the last energy state, electron density plots for both *Af*LPMO16 and *Af*LPMO_7EXK show that one side of the copper atom is free; there is no electron density. This is a clear indication of the pre-substrate binding state. The plot of the energy gap between HOMO and LUMO states of *Af*LPMO16 is 0.87eV, and the energy gap is 0.52eV for *Af*LPMO_7EXK (Fig. 4). Other mutant LPMOs show a higher energy gap.

**Figure 3.**
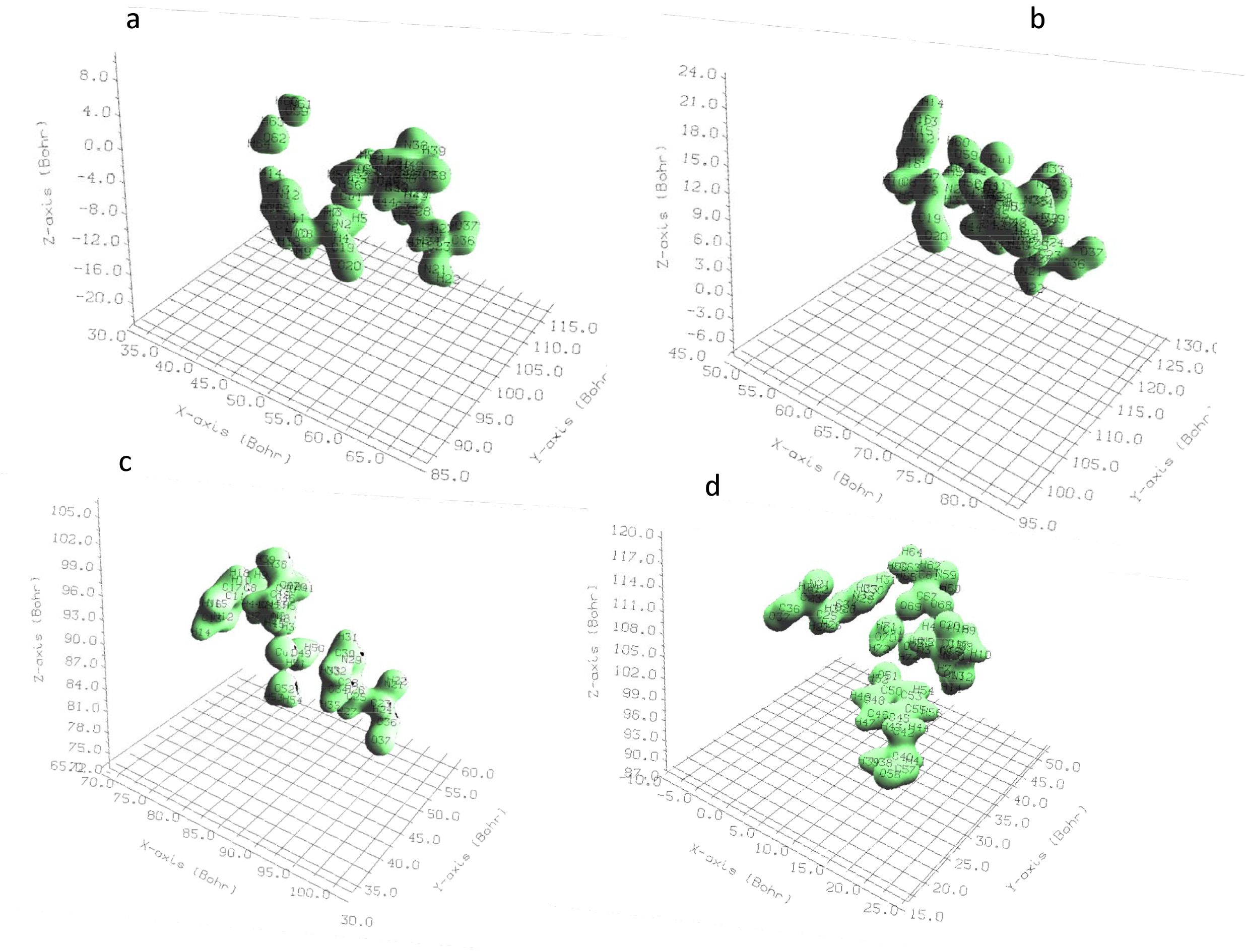
(a) Plot of electron density of active centre of 1^st^ minimize energy state of *Af*LPMO16 with the radius of 5Å. (b) Plot of electron density of active centre of last frame energy state of *Af*LPMO16 with the radius of 5Å. (c) Plot of electron density of active centre of 1^st^ minimize energy state of *Af*LPMO_7EXK with the radius of 5Å. (d) Plot of electron density of active centre of 1^st^ last frame energy state of *Af*LPMO_7EXK with the radius of 5Å.

**Figure 4.**
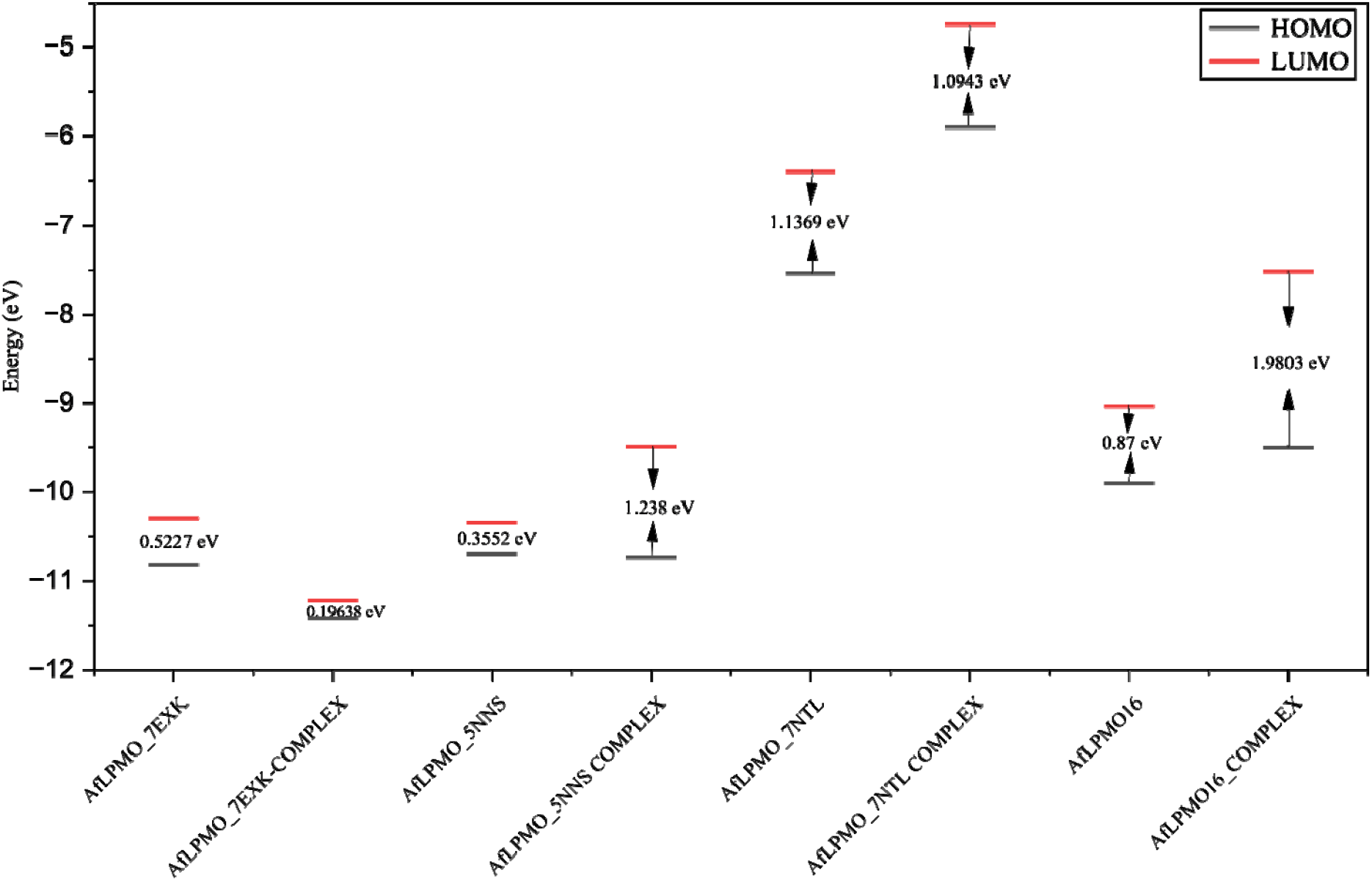
Plot of energy gap (electron volt) between highest occupied molecular orbital (HOMO) and lowest unoccupied molecular orbital (LUMO) energy of *Af*LPMO16 and chimeric proteins *Af*LPMO_5NNS, *Af*LPMO_7NTL and *Af*LPMO_7EXK with substrate bound and unbound state.

### Expression and Purification of *Af*LPMO16 and Its Mutants

*Af*LPMO16 and its L2 loop replaced mutants have nearly equal molecular weight with a maximum difference of 2kDa. All these mutants are purified, and SDS-PAGE analysis of the purified proteins suggested the proper molecular weight of the purified proteins (Fig. 5). Further, we sequenced the protein to check its native N-terminal expression, which also suggests the protein was expressed with its native N-terminal. Western blot analysis with anti-His antibody also confirms its proper expression (Fig. S2).

**Figure 5.**
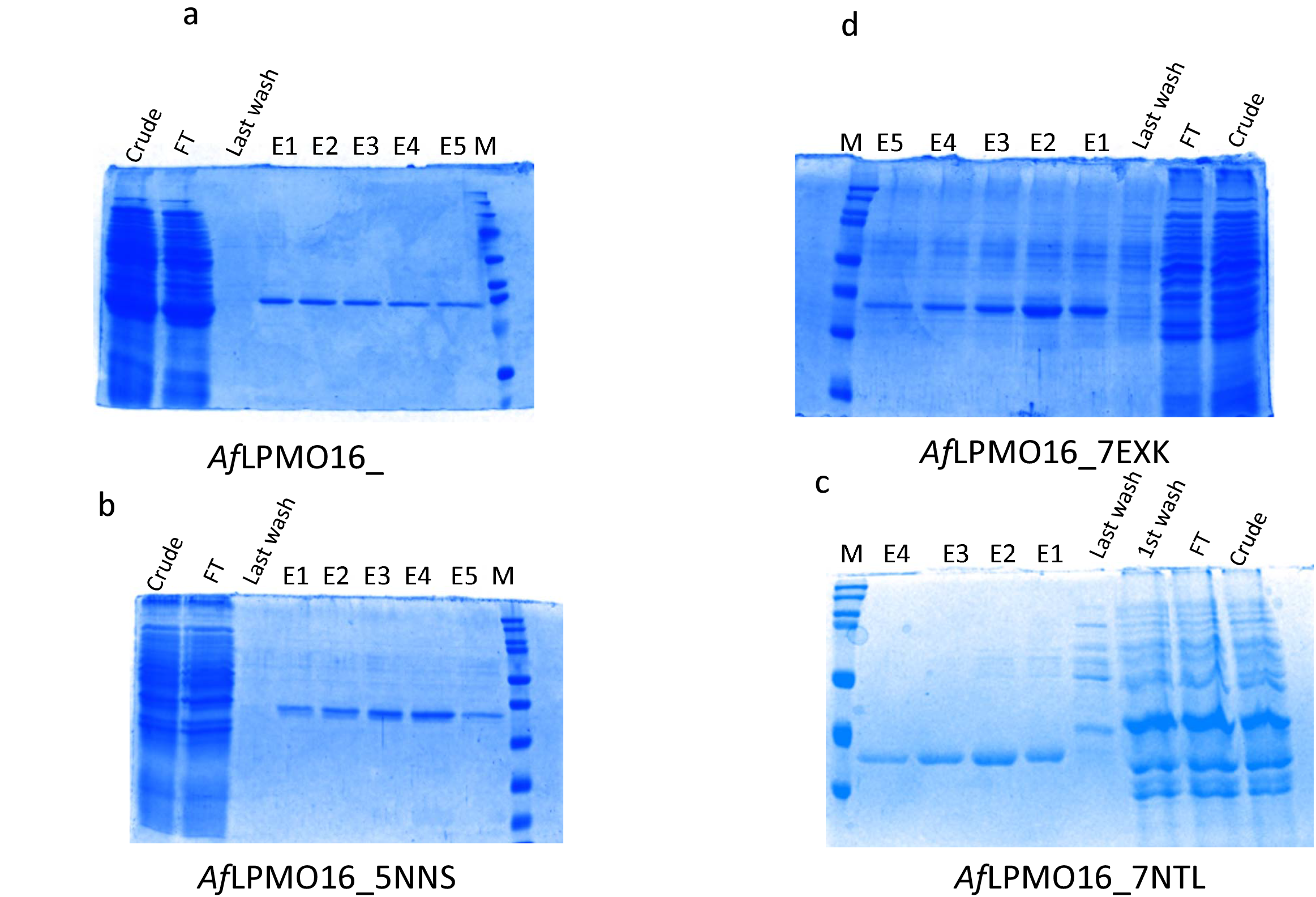
SDS-PAGE analysis of the (a) *Af*LPMO16 and chimeric proteins (b) *Af*LPMO_5NNS, (c) *Af*LPMO_7NTL and (d) *Af*LPMO_7EXK

### Enzyme activity

We performed three different assays to check activity and stability of *Af*LPMO16 and mutants: *Af*LPMO_5NNS, *Af*LPMO_7NTL and *Af*LPMO_7EXK. The Amplex-red assay measures the amount of hydrogen peroxide released by LPMO without a cellulosic substrate. Here, we performed the assay, and interestingly, we found that *Af*LPMO_5NNS released 51.2 μM of H_2_O_2_, which is the highest compared to wild-type *Af*LPMO16 and other mutants (Fig. 6a). *Af*LPMO16 and *Af*LPMO_7EXK released almost the same amount of hydrogen peroxide, 44.5μM and 44μM, respectively. The lowest amount of H_2_O_2_ released by *Af*LPMO_7NTL (29.6μM).

**Figure 6.**
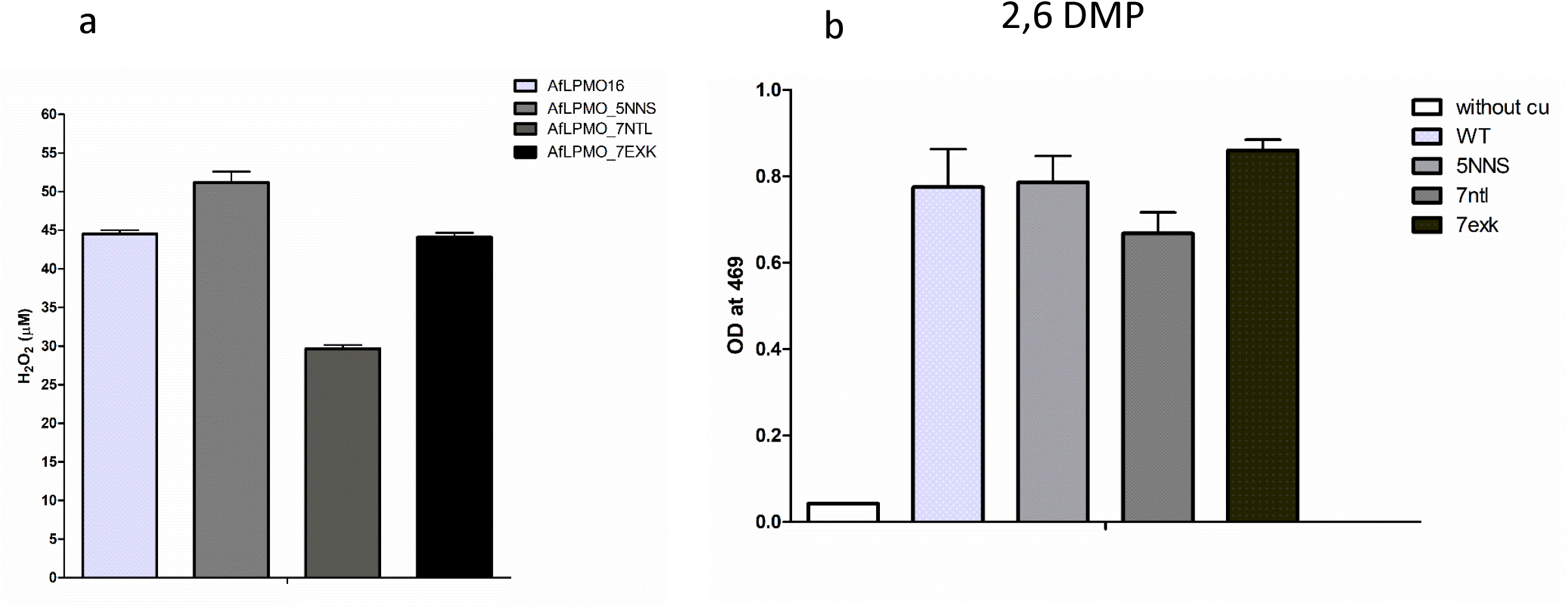
Enzyme assays: (a) Amlpex-red assay of *Af*LPMO16 and chimeric proteins *Af*LPMO_5NNS, *Af*LPMO_7NTL and *Af*LPMO_7EXK; in Y-axis the amount of hydrogen peroxide released by enzymes are measured. (b) Measurement of coloured product 1—coerrulignone produced by *Af*LPMO16 and chimeric proteins *Af*LPMO_5NNS, *Af*LPMO_7NTL and *Af*LPMO_7EXK; in Y-axis the OD is ploted at 469nm.

The 2,6-DMP is a non-physiological substrate to check the activity of LPMO. It measures the product 1-coerrulignone produced by the enzyme. Here in this assay, the results strongly suggest mutant *Af*LPMO_7EXK has the highest activity, followed by *Af*LPMO16, *Af*LPMO_5NNS, and *Af*LPMO_7NTL (Fig. 6b).

### Structural modifications of chimeric proteins

Biophysical studies like circular dichroism can reveal the secondary structural alteration of the mutant proteins compared to the wild type. The circular dichroism data of all these proteins indicated a β-sheet-dominated structure and very little α-helical content (Table 1). *Af*LPMO16 and *Af*LPMO_7EXK show almost similar α-helical content, 3.4% and 3.1% respectively. Whereas *Af*LPMO_7EXK showed a decrease in β-strand. On the other hand, did not show helical content, though the β-strand percentage is increased for these two mutants. Both of them have the 43.8% and 48.6% β-strand, respectively.

**Table 1.**

| Enzyme | Helix | β-strand | Turn | Other |
| --- | --- | --- | --- | --- |
| AfLPMO16 | 3.4% | 38.7% | 15.7% | 42.2% |
| AfLPMO_5NNS | 0% | 43.8% | 20.7% | 35.4% |
| AfLPMO_7NTL | 0% | 48.6% | 18.9% | 32.5% |
| AfLPMO_7EXK | 3.1% | 25.1% | 12.5% | 59.3% |

### Synergistic effects of the wild type and its mutant

Synergism of LPMO with cellulase enzyme is a crucial characteristic of the enzyme. To measure the synergism level of LPMO, the enzyme must be added with cellulase on a cellulosic substrate. Here, over a 48-hour time span, we measured the glucose release by the combined enzymatic reaction. The initial 6 hours of synergism of the *Af*LPMO16 are 2-fold. Where the *Af*LPMO_7EXK showed more than 2-fold synergism (Fig. 7). The synergistic effect is lowest for *Af*LPMO_7NTL almost 15% enhancement in glucose release. Throughout the 48-hour span, the *Af*LPMO_7EXK showed the highest activity, which is more than two-fold which strongly indicating its enhanced activity and stability.

**Figure 7.**
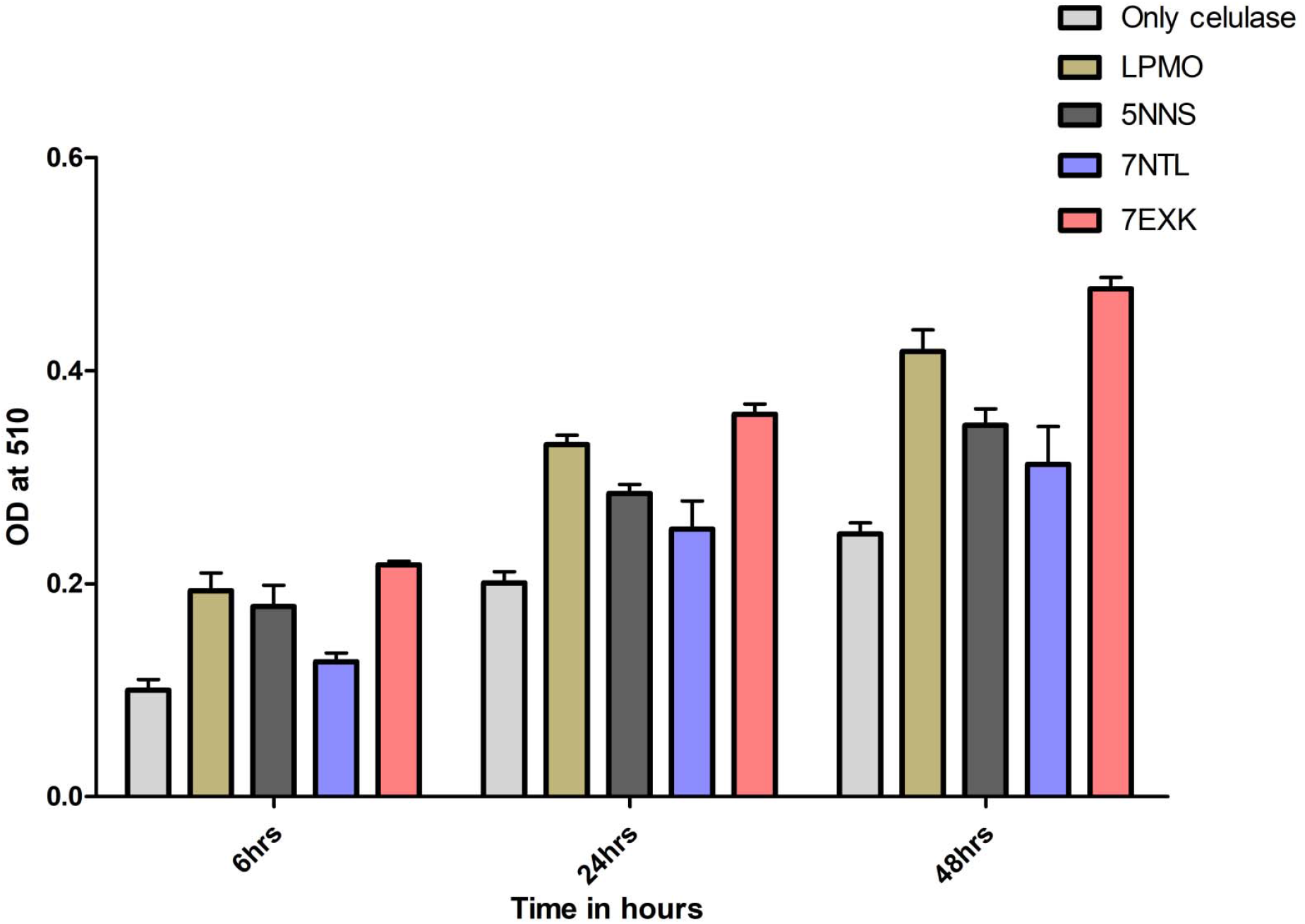
The synergism effect of *Af*LPMO16 and chimeric proteins *Af*LPMO_5NNS, *Af*LPMO_7NTL and *Af*LPMO_7EXK.

## Discussion

The enzymatic activity of the *Cs*LPMO9 (PDB: 7EXK) from the AA9 family is reported to be the highest activity using the 2,6-DMP as a substrate. Also, the enzyme is reported to show a 3.2-fold increase in yield due to synergism. So, this enzyme must be considered as a highly active LPMO among other reported LPMOs. The core idea of loop engineering is to determine which part of this enzyme is responsible for its higher activity. Whether it is the catalytic centre or the substrate binding loops, or the other non-catalytic part of the enzyme. The catalytic core of LPMO is almost similar; the difference is in the substrate binding loop or in the non-catalytic part of the LPMO. The substrate binding loop is the most obvious target that is completely or partially responsible for the enzymatic activity. That is why we targeted the L2 loop to replace it with the native one, and checked how the activity changes. The first challenge was to generate a stable chimeric protein because we are replacing a chunk of amino acids. We have successfully created these chimeric enzymes.

Then our aim is to validate the loop replacement engineering; in doing so, we have successfully engineered the L2 of LPMO, specifically the AA16 family LPMO from *A*.*fumigatus*. The computational studies, along with the expression validation, prove the formation of a stable chimeric protein with proper enzymatic activity. Further, the circular dichroism data suggest a significant amount of secondary structural changes in *Af*LPMO_5NNS and *Af*LPMO_7NTL decreased the helical content, that probably responsible for substrate binding in the loop region. However, the increment in β-strand percentage may stabilise the protein but perturb the substrate binding site. This could result in a decrease in enzymatic activity. The CD data confirms the proper structural prediction by the computational study. Further, the computational study also predicted the bond formation between the copper ion and Histidine residues in the active site of the enzyme, which also confirms the structural integrity. Quantum computational study revealed the electronic energy state, the energy gap between the highest occupied molecular orbital (HOMO) and the lowest unoccupied molecular orbital (LUMO). The gap between HOMO and LUMO for *Af*LPMO_7EXK is decreased to 0.52eV, which is less than *Af*LPMO16. That signifies the higher reactivity of the mutant enzyme *Af*LPMO_7EXK. This energy gap is increased for the other two mutants, which clearly indicates the lower reactivity of *Af*LPMO_5NNS and *Af*LPMO_7NTL. Even the substrate-bound *Af*LPMO_7EXK showed a lower energy gap of 0.193 eV, which strongly suggests the higher reactivity of the enzyme. Molecular dynamics study predicts the stability of the mutant proteins; further, the RMSF plot of *Af*LPMO_7EXK suggests higher flexibility of the mutant enzyme, which is also a signature of higher activity.

Expression of the chimeric protein with the native N-terminal is crucial for the enzymatic activity of LPMO. We confirmed it by protein sequencing (MALDI-TOF MS/MS) and proceeded with the enzymatic activity. First, we checked the peroxigenase activity or the uncoupling reaction (34). The H_2_O_2_ release by the mutant enzyme *Af*LPMO_5NNS is 51.2 μM, which is the highest among the mutants and the wild-type enzyme, suggesting a higher side reaction of this mutant. This higher amount of H_2_O_2_ could lead to self-inactivation (36). *Af*LPMO16 and *Af*LPMO_7EXK showed the same H_2_O_2_ release, which signifies similar side reactions and stability (37, 38). The *Af*LPMO_7NTL released the lowest amount of H_2_O_2_ due to its lower activity. This result is significant and has a close association with the enzyme assays. The 2,6-DMP assay clearly suggests the higher activity of *Af*LPMO_7EXK, which strongly indicates the L2 loop controls the activity. The lower activity of *Af*LPMO_7NTL also supports the Amplex-red assay data (34). Lastly, the synergistic effect of *Af*LPMO_7EXK is highest and throughout the 48-hour time period, it is stable. That again confirms the altered L2 loop is governing the complete enzymatic activity. Overall, this study confirms three crucial points: first, that the loop-altered enzyme can be stable and enzymatically active. Second, the L2 almost governs the enzymatic activity in the case of LPMO. The difference between enzymatic activity, synergism, and side reactions of LPMO and enzyme stability (preventing self-inactivation) could be controlled by only the L2 loop.

## Conclusion

In the concluding remark, this study proves that the L2 loop is controlling the overall reaction rate, and this loop alteration method is very useful and new to the LPMO chemistry field. We first report the successful L2 loop engineering for LPMO. But there few questions that need to be answered in future studies. Does the loop completely control the regioselectivity? Whether only the loop alteration is sufficient to change the substrate specificity. What are the structural changes by the replacement of the L2 loop?

## Acknowledgment

MH is thankful to the DBT (Govt. of India) for the fellowship (Grant No. BT/PR41285/PBD/26/804/2020). SD is thankful to DST INSPIRE (Govt. of India) for the research fellowship.

## Author contributions

MH conceptualised and articulated. MH, SD, NB, and SR performed the experiments. MH and SD wrote the manuscript. SSM and BK reviewed and edited the manuscript. All the authors have reviewed and approved the final manuscript.

